# The distribution of the hemlock woolly adelgid in Canada

**DOI:** 10.1101/2024.07.18.604084

**Authors:** Chris J K MacQuarrie, Meghan Gray, Erin Bullas-Appleton, Troy Kimoto, Nicole Mielewczyk, Ron Neville, Jeffrey B Ogden, Jeffrey G Fidgen, Jean J. Turgeon

## Abstract

The hemlock woolly adelgid, *Adelges tsugae* Annand (Hemiptera: Adelgidae) has distinct native and invasive populations in Canada. On Canada’s west coast the adelgid is a native insect feeding on western hemlock, *Tsuga heterophylla* (Raf.) Sarg. and mountain hemlock, *Tsuga mertensiana* (Bong.) Carrière (Pinaceae). In eastern Canada, the adelgid is an invasive species that attacks and kills eastern hemlock, *Tsuga canadensis* (L.) Carrière (Pinaceae). We obtained all records of *A. tsugae* in institutional and public databases and develop updated range maps, phenologies and dispersal estimates for the species in British Columbia and eastern Canada In British Columbia *A. tsugae*’s distribution is centred around the lower mainland and on Vancouver Island but with populations in the interior and along the Pacific coast that have been poorly explored and which could be sources of biological control agents to manage invasive populations in the east. In eastern Canada, the adelgid has invaded all of southern Nova Scotia, portions of the Niagara region in Ontario as far west as Hamilton, and at least one site on the north shore of Lake Ontario. No populations have been found in New Brunswick, Quebec or Prince Edward Island. Finaly, we estimated the rate of spread in Nova Scotia at 12.6 ± 8.2 to 20.5 ± 27.21 km/year.

## Introduction

Hemlock trees, *Tsuga* spp. (Endlicher) Carrière (Pinaceae), are an important component of late successional forests in eastern and western Canada. Three species of hemlock are native to Canada: *Tsuga heterophylla* (Raf.) Sarg. (western hemlock), and *Tsuga mertensiana* (Bong.) Carrière (mountain hemlock) with sympatric distributions in British Columbia, and *Tsuga canadensis* (L.) Carrière (eastern hemlock), which occurs throughout southern Ontario, southern Quebec and the Maritime provinces (Figs. 1,2,3). Hemlocks are long-lived, shade-tolerant tree species that typically grow in cool, moist environments and on a wide variety of soil types and origins. In British Columbia, *T. heterophylla* grows in association with many other tree species and understory shrubs and is often an indicator of climax or near-climax communities. It can be found from sea level to >2000 m above sea level (asl) whereas *T. mertensiana* grows in the subalpine zone in British Columbia between 900-1,800 m asl and grows best in sheltered, mixed-species stands with northern exposure. In the east, *T. canadensis* typically grows on acidic, moist soils with good drainage found from sea level to 730 m asl and often has very little understory under well developed stands and is where it is a major component of four forest types, either as the dominant species or in association with white pine, *Pinus strobus* Linnaeus (Pinaceae); yellow birch, *Betula alleghaniensis* Britt. (Betulaceae); or yellow poplar, *Liriodendron tulipifera* Linnaeus (Magnoliaceae) as well as American beech, *Fagus grandifolia* Ehrhart (Fagaceae); red spruce, *Picea rubens* Sargent (Pinaceae); and sugar maple, *Acer saccharum* Marshall (Sapindaceae) on upland sites (Summarized from: Godman and Lancaster 1990; Means 1990; Packee 1990). In eastern Canada the abundance of eastern hemlock has been reduced by as much as 80% since European colonization and now comprise < 4% of forest cover in its previous range (Loo and Ives 2003; Emilson et al. 2018). This is mostly due to conversion of land to agriculture and other uses, and the preference for early-successional tree species in commercial forestry. What hemlock remains has high ecological and social value in eastern Canada (Parker et al. 2023). In western Canada hemlock retains much of its original range and provides ecological benefits in riparian ecosystems, but the species is also a significant component of coastal and subalpine forests where it is a commercially important species.

**Figure 1.**
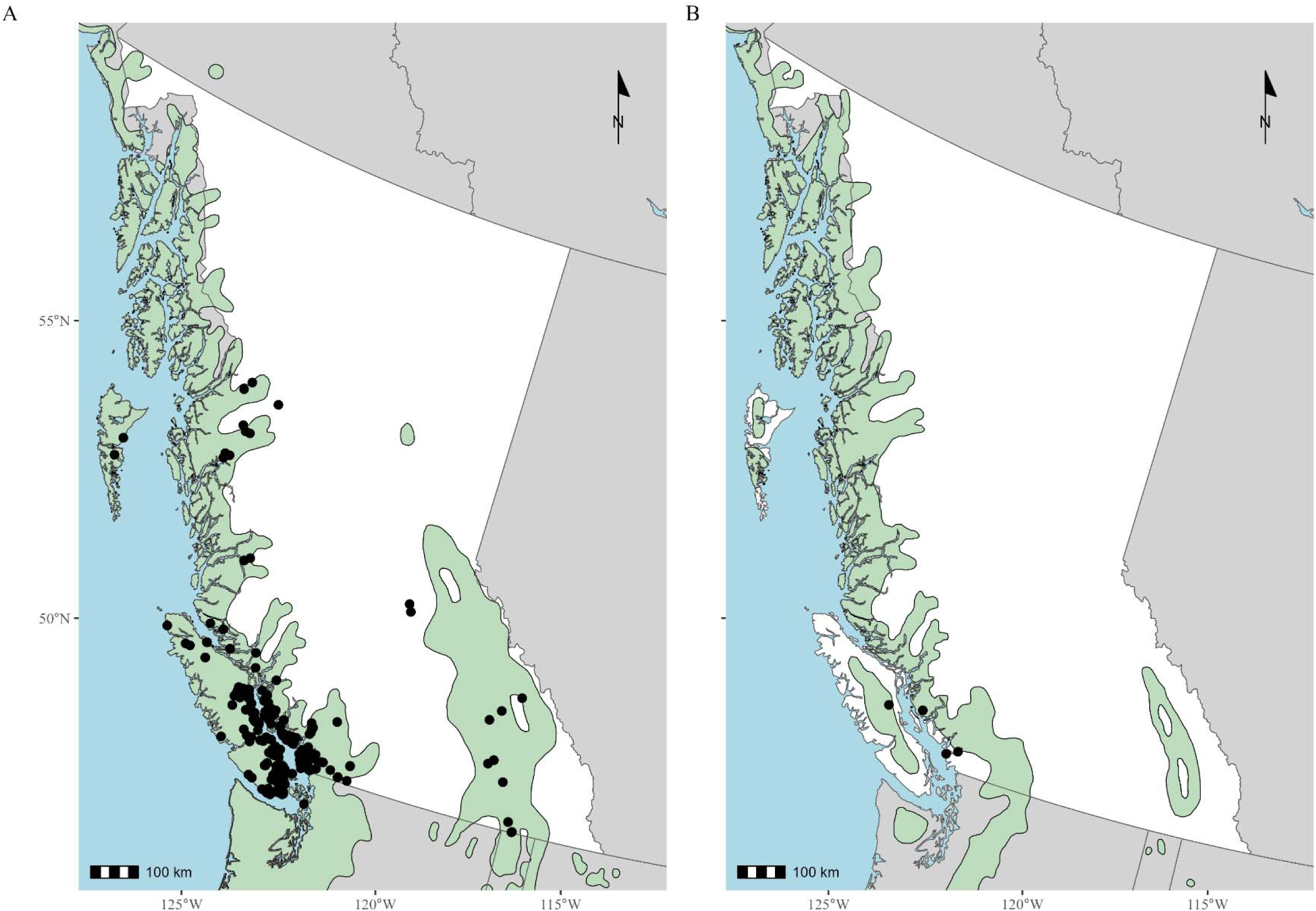
Recorded observations (circles) of *Adelges tsugae* on *Tsuga heterophylla* (A) and *Tsuga mertensiana* (B) in British Columbia, Canada. Green shaded areas show the distribution of each host tree species (Fryer 2018). Data sources for *A. tsugae* observations are given in the text; map data from Natural Earth; map A includes all observations where the host was recorded as ‘*Tsuga*’.

**Figure 2.**
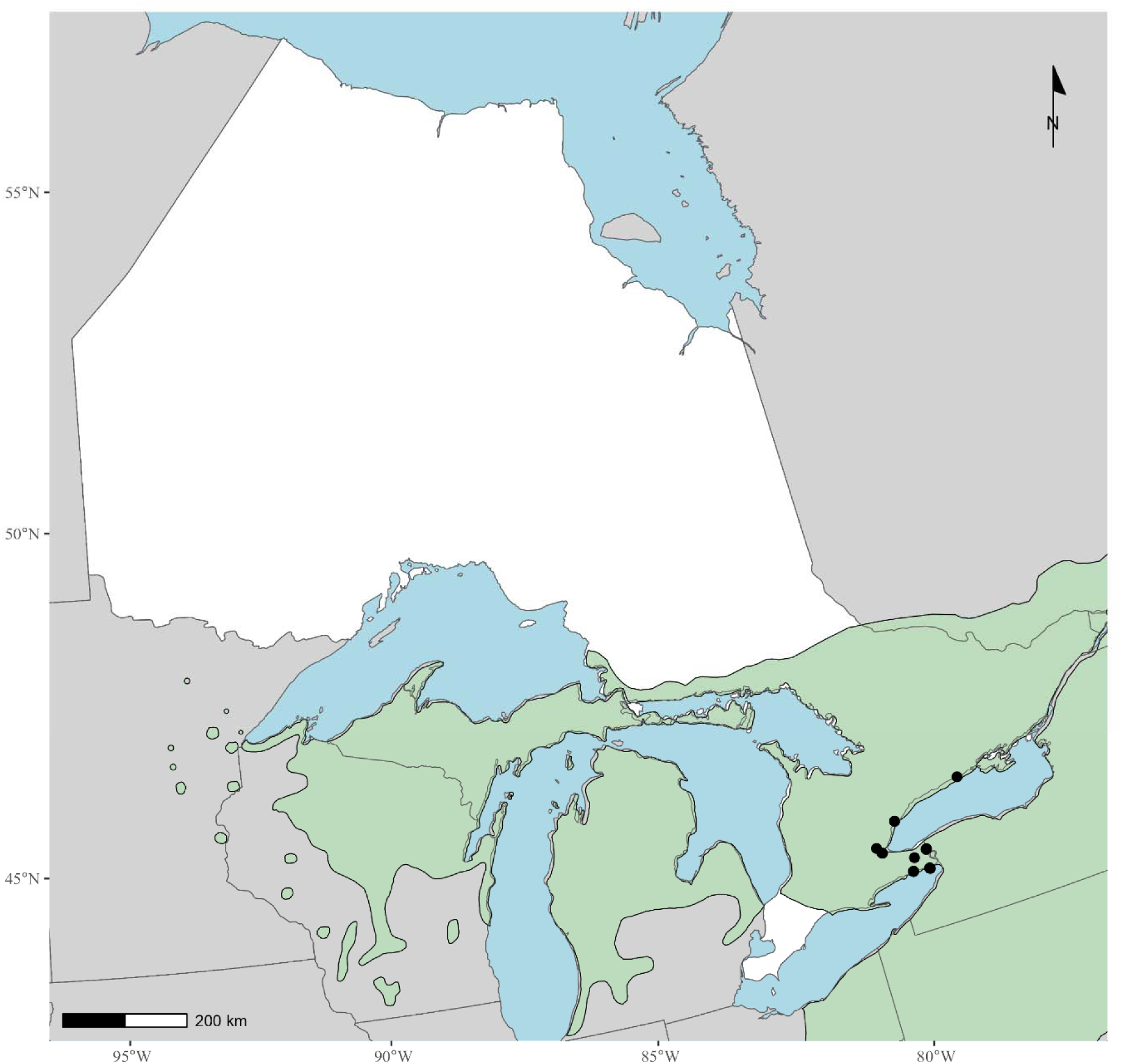
Recorded observations (circles) of *Adelges tsugae* as of 2023 in Ontario, Canada and the range (green) of *Tsuga canadensis*. Data sources for *A. tsugae* observations are given in the text; distribution of *T. canadensis* from Fryer (2018); map data from Natural Earth.

**Figure 3.**
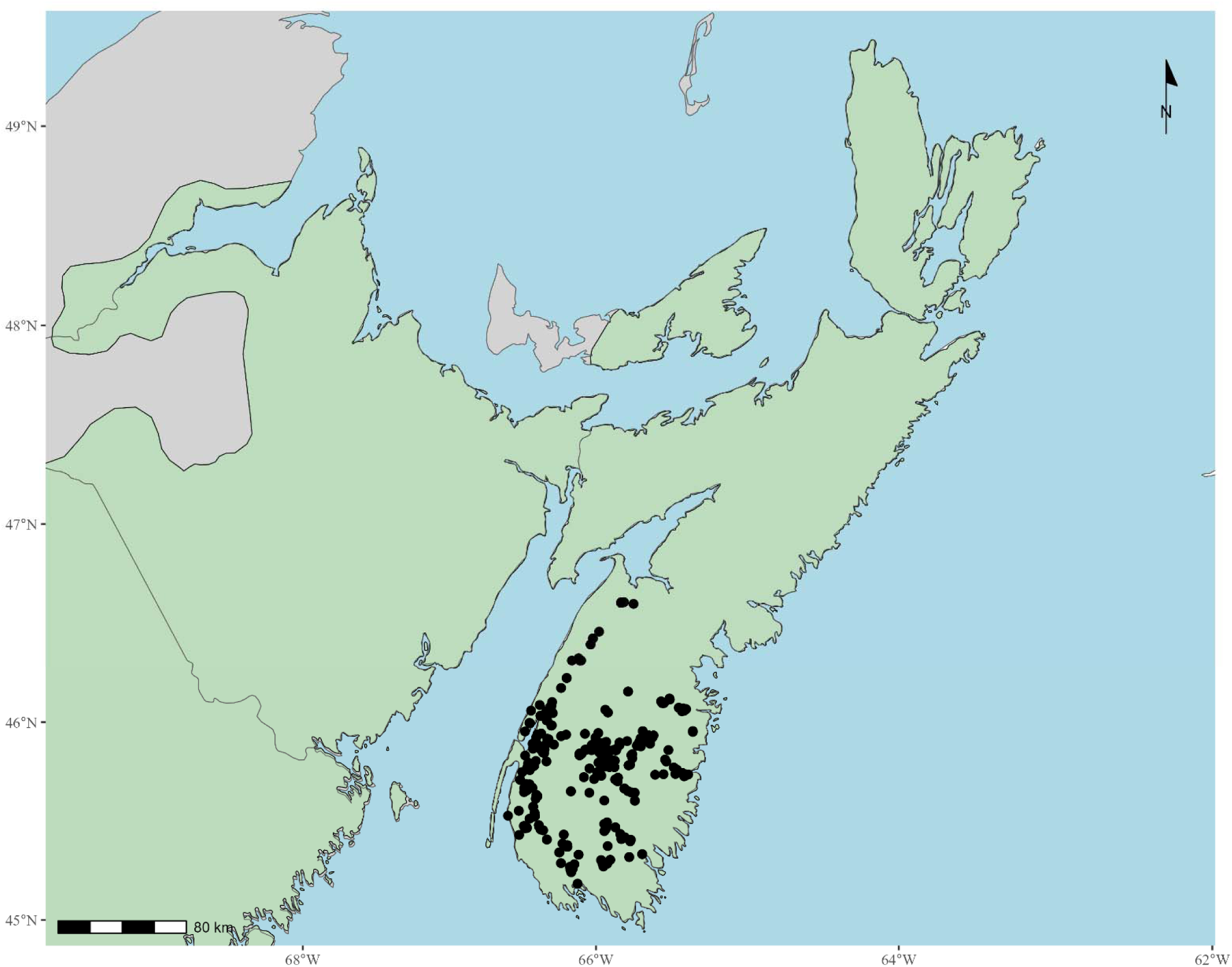
Recorded observations (circles) of *Adelges tsugae* as of 2023 in Nova Scotia, Canada and the range (green) of *Tsuga canadensis*. Data sources for *A. tsugae* observations are given in the text; distribution of *T. canadensis* from Fryer (2018); map data from Natural Earth.

In the eastern United States of America *T. canadensis* and *Tsuga caroliniana* Engelm. (Carolina hemlock) have been under threat from *Adelges tsugae* Annand (Hemiptera: Adelgidae), the hemlock woolly adelgid, for at least the past 70 years. This invasive species was first discovered at a private arboretum in Richmond, Virginia in the 1950s and has been slowly spreading in the eastern US range of hemlock since the 1960s. The insect was likely introduced from Japan (Havill et al. 2006; Havill et al. 2016a), possibly on infested Japanese hemlocks intended for horticultural planting. In the late 1980s, *A. tsugae* gained prominence as an significant invasive species when hemlock mortality was observed in the mid-Atlantic states (McClure 1987; McClure 1989). Spread of the insect has been slow, estimated at 5 to 20 Km per year (Morin et al. 2009; Turner et al. 2011; Fitzpatrick et al. 2012; Goldstein et al. 2019), and likely facilitated though long distance spread by migrating birds (McClure 1990; Russo et al. 2016; Russo et al. 2019), the movement of infested wood or plants (e.g., North American Plant Protection Organization 2012), and local spread by wind and mammals (McClure 1990; Turner et al. 2011). The northward spread of *A. tsugae* was thought to be constrained by cold winter temperatures limiting the insect’s survival (Parker et al., 1998; Parker et al., 1999; Skinner et al., 2003), but recent research showed that its overwintering cold tolerance could facilitate northward spread (Paradis et al. 2008; Elkinton et al. 2017; Lombardo and Elkinton 2017). This hypothesis was supported in by detections of infestations in coastal Maine, Vermont, upstate New York, Ontario, and Nova Scotia (Canadian Food Inspection Agency 2012; 2017; Hemlock Woolly Adelgid National Initiative 2024; Nova Scotia Department of Lands & Forestry and Nova Scotia Department of Environment and Climate Change 2024). Recent modelling, based on overwintering and survival data from populations in the eastern United States of America suggests that under both present and future climate some, if not all, of the range of eastern hemlock in Canada is at risk of invasion by hemlock woolly adelgid (McAvoy et al. 2017; Ellison et al. 2018; Kantola et al. 2019; Cornelson et al. in press). In the eastern United States of America *T. canadensis* trees die within 4-15 years following the initial attack and their within-stand mortality can exceed 90% (McClure 1991; Orwig and Foster 1998; Eschtruth et al. 2013). *Adelges tsugae* is therefore thought to have the potential to have significant negative impacts on the remaining hemlock forests in eastern Canada (Emilson et al. 2018; Parker et al. 2023).

*Adelges tsugae*, though invasive in eastern North America, is an endemic, native species in western North America. The western population of *A. tsugae* likely originates from natural spread by the insect during the time North America and Asia were connected (Havill et al. 2006; Havill et al. 2016a) and not the result of recent invasion. This long association has resulted in co-evolution between *A. tsugae* and a suite of natural enemies that appears to regulate populations of the insect (Crandall et al. 2022). A similar regulatory mechanism is not observed in eastern North American populations and is likely responsible for *A. tsugae*’s success as an invasive species (Wallace and Hain 2000; Mayfield III et al. 2023). Natural enemies of *A. tsugae* from western North America have therefore been released as biological control agents in eastern North America, as well as natural enemies from Asia (Mayfield III et al. 2023).

The range of *A. tsugae* in western North America is not well documented but is assumed to be roughly the same as the range of western and mountain hemlock. Kantola et al. (2019) recently accumulated occurrence records for this species from North America and Asia to inform the development of a species distribution model. Most of the observations used by Kantola et al. (2019) for western North America appear to originate from the iNaturalist database (www.iNaturalist.com) augmented with those from Havill et al. (2016a) which only contain four records from British Columbia, all from the lower mainland or Vancouver Island. A fifth record from the region indicates that *A. tsugae* was also found on Prince of Wales Island in the Alaskan panhandle. The records for western Canada used by Kantola et al. (2019) are incomplete, however, as hemlock woolly adelgid is known to inhabit other areas of the province. For example, British Columbia has been the source of at least one biological control agent used in eastern North America that originated from Vancouver Island (Zilahi-Balogh et al. 2003; Zilahi-Balogh et al. 2012). Therefore, there is a need to document observations of *A. tsugae* in western Canada to inform research efforts (e.g., Kantola et al. 2019), including prospecting for potential biological control agents in its native range.

Similarly, since 2012 there have been a series of introductions and detections of *A. tsugae* in eastern Canada, most notably in southern Nova Scotia in 2017. The data on the location of pest detections are critical to understand the potential for range expansion and assessing the risk of *A. tsugae* to *T. canadensis* in eastern Canada. These data would also allow us to validate predictions from models for climatic suitability developed using populations from the eastern United States of America. Information about these detections have not been consolidated and only basic information on the detections are contained in annual reports (e.g., Canadian Food Inspection Agency 2012), and no data on spread is contained in these reports. It is also important to consolidate this information because in all of eastern Canada *A. tsugae* is considered a pest regulated by the Canadian Food Inspection Agency (CFIA) under the authorities of the Plant Protection Act and Regulations (1990). As part of this regulation the CFIA identifies the specific areas of Canada, and the United States of America, where *A. tsugae* is considered to be established and outlines the movement requirements for *A. tsugae* ’s host material to and from these regulated areas. The CFIA also has an obligation to international trading partners to conduct surveillance activities for *A. tsugae* within unregulated areas of Canada to provide data in support of CFIA’s declarations of pest freedom for commodities originating from areas where the pest is not known to occur. Once a new find of *A. tsugae* is identified within an area of Canada where the pest was not known to exist, the CFIA is required to take regulatory action to mitigate the immediate risk of artificial spread. At the same time, the CFIA conducts delimitation surveys to assess the overall size and severity of the infestation and determine response options. The response may involve implementing controls to eradicate the population or the establishment of larger regulated areas to mitigate the risk of spread from the area.

Herein, we accumulate all records of *A. tsugae* in Canada and use these data to clarify the present distribution in British Columbia and understand the spread of the insect in eastern Canada. We show that the occurrences of this adelgid in British Columbia suggest that many areas are under-explored for *A. tsugae* and it’s associated natural enemies. We also show that contemporary invasions in eastern Canada are spreading at rates consistent with those observed in the eastern United States of America.

## Methods

We used published and unpublished records of collections of *A. tsugae* from a variety of sources to determine its range in Canada.

Specimen data records for *A. tsugae* from western Canada were extracted from institutional and publicly available databases. We obtained collection records of the Canadian Forest Insect and Disease Survey held within the Forest Invasive Alien Species Document library (Natural Resources Canada Canadian Forest Service 2023). These records are observations of insects or damage reported as part of annual forest health surveys done in Canadian forests by the Canadian Forest Service from 1936-1994. We also included records in the Forest Insect and Disease Survey database from before 1936 that originate from earlier surveys (Van Sickle et al. 2001). Records from the Forest Insect and Disease Survey included host records so we included that information as well. We supplemented the Forest Insect and Disease Survey data with records of *A. tsugae* for Canada in the Global Biodiversity Information Facility which includes records of observations from iNaturalist, and specimen data records from arthropod collections including the Canadian National Collection of Insects, Arachnids and Nematodes and the Yale Pebody Museum (Global Biodiversity Information Facility 2023). These data lacked host records. We also included records from: the Barcode of Life database (Ratnasingham and Hebert 2007); the collections of Natural Resources Canada Canadian Forest Service’s Pacific Forestry Centre; Havill et al. (2016a) which we obtained from Havill et al. (2016b), and observations made by two of us (JGF, TK) between 2019 and 2023 in British Columbia during explorations for *A. tsugae* biological control agents by the Canadian Forest Service, CFIA and Cornell University. Inspection of the resulting combined database suggested there was a small amount of duplication in records from institutional collections, The Barcode of Life database, and Havill et al. (2016b). In particular, for specimens collected on southern Vancouver Island and the Vancouver region. This means a small number of records from this region may be duplicated in our database. We also note that early records in the Forest Insect and Disease Survey database for British Columbia were often imprecise as to the geographic location of the observations. The Forest Insect and Disease Survey records report this geographic accuracy, so we limited our examination to just those records where the accuracy was < 20 Km.

Records for eastern Canada were obtained by the authors from CFIA survey and reporting database (provided by RN) and Nova Scotia Department of Natural Resources and Renewables (provided by JBO). Once an area has been regulated under an Infested Places Order by the CFIA the agency ceases its intense surveys within that area. So we augmented the CFIA data with iNaturalist records in GBIF (GBIF.org, 2023). There were no records for eastern Canada in any of the other databases we inspected.

Data from the CFIA include the first detection records for *A. tsugae* in eastern Canada. The year and location of each of these first detections are contained in annual survey reports of the CFIA (2012-; 2023). For each of these first detection records we provide a description of how these detections were made based on contemporary records, reports to the CFIA, and the authors’ own observations that were not included in the annual survey reports. We also evaluated the location and yearly distribution of CFIA survey sites for *A. tsugae* in eastern Canada over the same period to evaluate survey effort.

We evaluated the temporal trend in hemlock woolly adelgid collections using data associated with each collection record. For records from British Columbia we determined the altitude for each specimen by cross referencing the recorded locations to a digital elevation model for the province.

For records from Nova Scotia we examined the rate of spread for the invasive population from its initial detection location in 2017. We used two methods to evaluate range expansion in Nova Scotia. For the first method we used all records from the province, determined the geographic centre of the *A. tsugae* population in each year (the centroid) and determined the distance between centroids from consecutive years. These records included those from CFIA inspection data and records from Global Biodiversity Information Facility. From these values we computed the mean change in the position of the population’s centroid since 2017. The second approach repeated this analysis but evaluated the northward spread of the population by examining the change in position of the most northern observation in the province. We report the mean and standard deviation of our estimates.

All mapping and analyses were done in the R statistical computing environment (R Core Team 2019), except for the determinization of altitudes which was done in ArcGIS. Data are available from the Barcode of Life database (Ratnasingham and Hebert 2007), the Forest Invasive Alien Species Document library (Natural Resources Canada Canadian Forest Service 2023), Global Biodiversity Information Facility (GBIF.org, 2023) and Havill et al. (2016b). Data from CFIA and Nova Scotia Department of Natural Resources and Renewables can be obtained by contacting those agencies.

The range of *T. heterophylla*, *T. mertensiana* and *T. canadensis* were mapped using resources from Fryer (2018). These maps are known to be somewhat imprecise with respect to the range of some tree species and they do not reflect the abundance of each species. Herein, the maps are used only to show the approximate area each tree species can be expected to be found.

## Results and Discussion

### British Columbia

The earliest definitive record of *A. tsugae* in Canada is in a report from May 1938 indicating that the species was active on *T. heterophylla* in Vancouver, British Columbia (Venables and Hopping 1938; Hopping 1939). Chystal (1916), however, reported an insect of very similar description to *A. tsugae* on *T. heterophylla* in Vancouver’s Stanley Park 22 years earlier (See Havill et al. 2006 for further discussion). Most observations of *A. tsugae* in British Columbia were made during the 1960s through the late 1980s - early 1990s (Fig. 4A). There are no apparent reasons for the gap from 1970 to ca. 1985 but the abrupt end of records in 1994 corresponds with the demise of FIDS and the devolution of forest health surveys to the Canadian provinces.

**Figure 4.**
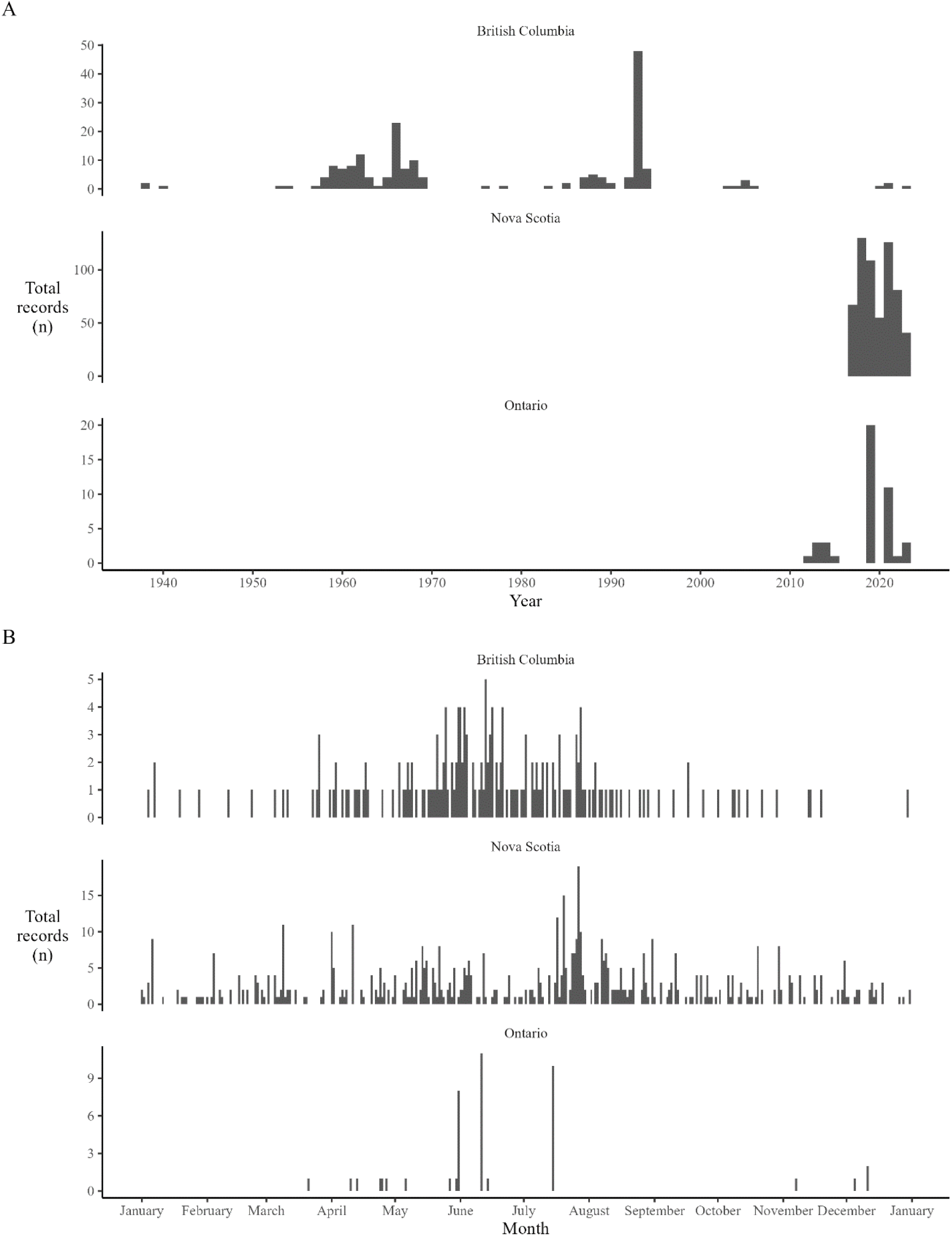
Historical timing (A) and within-year phenology (B) of *Adelges tsugae* observations in three Canadian provinces.

*Adelges tsugae* were collected year-round in British Columbia with most collections made during the summer months (Fig. 4B). The range of *A. tsugae* spans from 48.4 to 55.4 north latitude in British Columbia and from 0–1,800 m asl with most records coming from the coastal part of the province, which corresponds to British Columbia’s Coastal Western Hemlock and Mountain Hemlock Biogeoclimatic Zones (British Columbia Ministry of Forests - Research Branch 1997). Both *T. heterophylla* and *T. mertensiana* occur in this region and are generally restricted to the Pacific coast, although there are populations of both species in southeastern British Columbia. In general, the southern portion of *T. heterophylla*’s coastal range is well-surveyed for *A. tsugae* (Fig. 1A). In comparison, the more-northern coastal and interior forests appear to be relatively under-surveyed. The coastal region of British Columbia is difficult to access which has likely limited the investigations of *A. tsugae* in these areas. This may partly explain why, in that region, most observations appear to correspond to the location of roads that link the interior to the Pacific coast. Collections of *A. tsugae* have also been made on Haidi Gwaii (Fig. 1A) and on Alaska’s Prince of Wales Island (Havill et al. 2016b).

The eastern range of *T. heterophylla* in British Columbia does not appear to have been well surveyed for *A. tsugae* (Fig. 1A). In the southeastern part of the province *Tsuga* species are distributed through the Interior Cedar-Hemlock Biogeoclimatic zone and to a lesser extent in the Engelmann Spruce Sub-alpine Fir Biogeoclimatic zone (British Columbia Ministry of Forests - Research Branch 1996; 1998). The range of *T. heterophylla* in this region is disjunct from that along the Pacific coast, but is connected to populations in Oregon, Idaho and Montana. *Tsuga mertensiana* also occurs in this region but over a smaller area (Fig. 1B). There are about a dozen observations of *A. tsugae* in this part of the province, mostly in the Kootenay region of southeastern British Columbia (Fig. 1A), with no observations in the northern part of the range. Most observations are from the FIDS records, however, more recently HWA was found near Likely and on the southwest side of Quesnel Lake in the Cariboo region (TK, personal observation). The southern and lower elevation parts of this region in BC are climate zones 5 and 6 whereas the northern and higher elevation parts are zones 3 and 4 and may therefore not be a suitable for *A. tsugae*, based on climate conditions not being conducive to successful overwintering (e.g., Elkinton et al. 2017). The distribution of *Tsuga* spp., and thus also *A. tsugae*, may therefore be limited in this area. Even though *Tsuga* spp. can grow in wetter parts of the Cariboo region on the western side of the Cariboo mountains. The climate tolerances of *A. tsugae* in its native range in western North America have not been assessed. If *A. tsugae* does exist in this region it could host populations of natural enemies that may be better adapted to the regions at risk of invasion by *A. tsugae* in southern Ontario, southern Quebec, and the Maritimes (Figs.s 2 and 3). For example, a predator of *A. tsugae*, *Laricobius nigrinus* (Fender) (Coleoptera: Derodontidae), from the interior of Idaho were more cold tolerant than individuals of the same species from coastal Washington state (Mausel et al. 2011). These interior populations were therefore predicted to better match the climate in invaded regions of eastern North America where the predator is used as a biological control agent against *A. tsugae*. This would suggest that exploration for potential *A. tsugae* biological control agents for Canada may wish to consider this region of east-central British Columbia.

Most observations from the FIDS records report *A. tsugae* from *T. heterophylla* or from ‘*Tsuga* sp.’. There are, however, records of *A. tsugae* from *T. mertensiana* in 1966 and recent records of *A. tsugae* on *T. mertensiana* at the alpine ski resort on Mt. Washington on Vancouver Island, at Mt. Seymour Resort in Mt. Seymour Provincial Park near Vancouver, and at the University of British Columbia’s Botanical Garden in Vancouver and a record of *A. tsugae* on introduced *T. canadensis* at the Malcom Knapp Research Forest in Maple Ridge (TK, personal observation).The FIDS data also record *A. tsugae* being collected from Pacific silver fir, *Abies amabilis* (Dougl.) Forb (Pinaceae); Douglas-fir *Pseudotsuga menziesii* (Mirb.) Franco (Pinaceae); and western redcedar, *Thuja plicata* Donn (Cupressaceae). These observations from non-*Tsuga* spp. hosts represent a minute fraction of records, and of which three are from beat-sheet collections and could have therefore been contaminated by *A. tsugae* dislodged from neighboring trees (Natural Resources Canada Canadian Forest Service 2023). These specimens could also have been misidentified. For example, *A. amabilis* hosts *Adelges piceae* (Ratzeburg) (Hemiptera: Adelgidae) and *P. menziesii* hosts *Adelges cooleyi* (Gillette) (Hemiptera: Adelgidae), both of which could have been mistaken for *A. tsugae*.

### Alberta, Saskatchewan, and Manitoba

*Tsuga heterophylla* is recorded in the Rocky Mountains of western Alberta (Moss and Packer 1983) near the very edge of its range. Other members of the genus do not grow natively in the Prairie provinces and so there are no historical or contemporary records of the adelgid, and no surveys are conducted in any of those three provinces. *Tsuga* spp. are, however, likely planted in this region as ornamental trees and would be at risk of infestation if exposed to infested material imported from British Columbia, the Pacific northwest or eastern North America.

### Ontario

Multiple infestations of *A. tsugae* were found in Ontario between 2012 and 2024.

In 2012, an *A. tsugae* infestation was discovered on four landscape *T. canadensis* by an arborist at a private residence in the Etobicoke district of Toronto marking the first detection of *A. tsugae* in eastern Canada (Fig. 2). The infested trees were removed, and a visual delimitation survey was performed (Fig. 5A) by assessing all species of *Tsuga* within a 500 m radius of the infested site. In 2013, two additional infested trees were identified on neighboring properties, which were also removed and destroyed. Delimitation surveys were conducted for five years (2014-2019), and no additional infestations were detected in Etobicoke.

**Figure 5:**
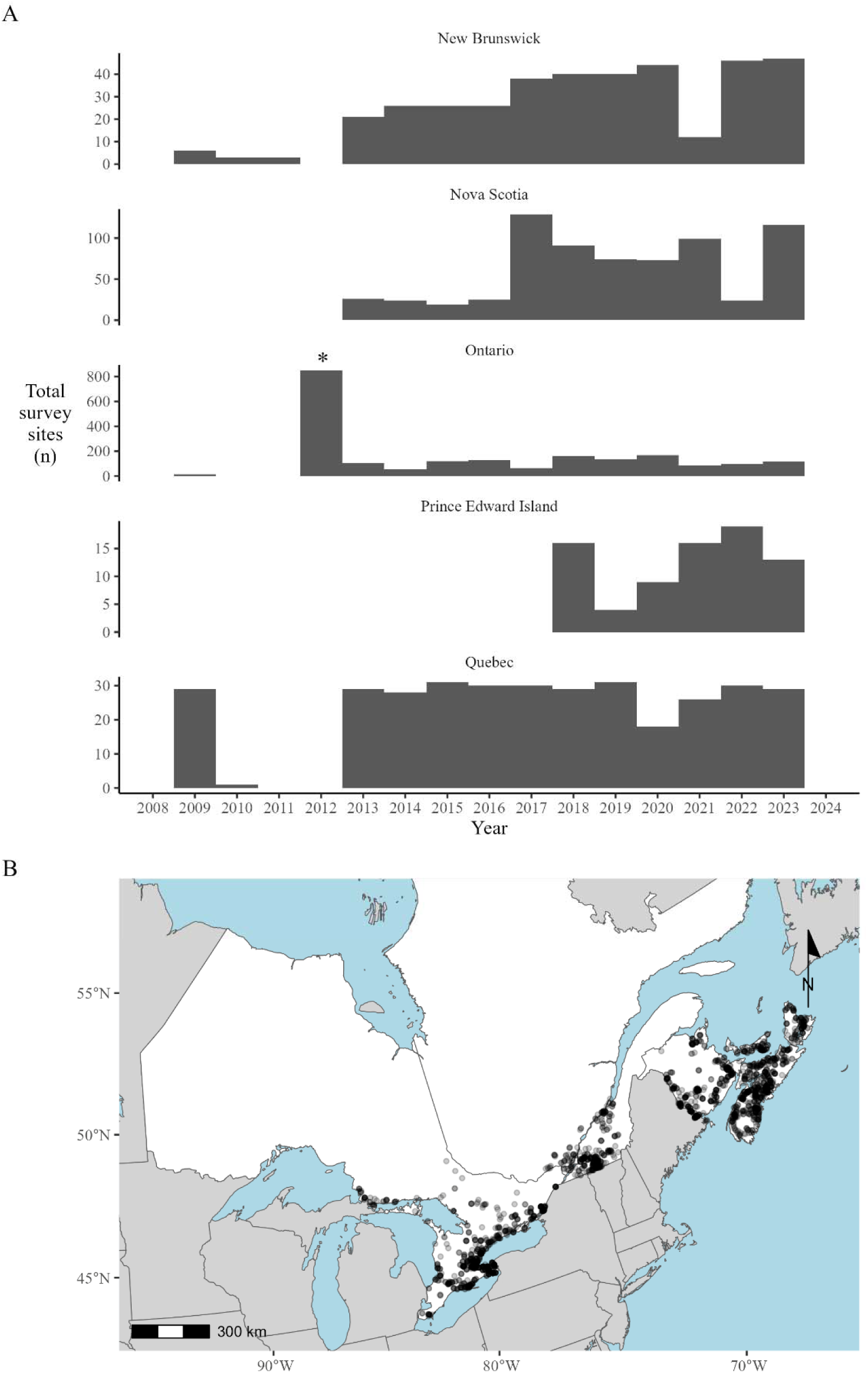
Number (A) and location (B) of Canadian Food Inspection Agency *Adelges tsugae* survey sites in Eastern Canada, 2008-2024. Regions with a higher density of survey points in (B) have darker shading. The large number of survey sites in Ontario in 2012 (* in A) reflect increased visual survey and delimitation efforts following the first detection in Canada, see text for more details.

In 2013, *A. tsugae* was found on one *T. canadensis* tree in the Niagara Gorge in the city of Niagara Falls during a routine detection survey targeting *A. tsugae* conducted by the CFIA and Canadian Forest Service. The infested tree was cut and burned on site and delimitation surveys using visual inspection, sticky card trapping (Fidgen et al. 2020), ball sampling (Fidgen et al. 2021) and branch sampling were implemented to determine the extent of the infestation but no additional *A. tsugae* infestations were discovered in Niagara Falls. Follow-up surveys were conducted starting in 2014 to verify the efficacy of the control efforts and to guide policy decisions. These surveys were conducted between November and June, the optimal time to visually assess hemlock trees for signs and symptoms of attack. As a result of these surveys additional infested trees were found in 2014 and 2015 in the Niagara Gorge. These trees were located adjacent to the single tree found in 2013. As in 2013, these trees were destroyed on site. Subsequent delimitation surveys were conducted annually, with no additional detections until 2019.

New infestations of *A. tsugae* have been in found in Ontario almost every year since 2019. In 2019, *A. tsugae* infestations were discovered near Wainfleet and in Niagara Falls during routine CFIA *A. tsugae* detection surveys. The 2019 detection in Niagara Falls was found on 11 trees (out of 92 examined) located downriver of the original detection in 2013 and outside the original delimitation zone. Compared to the single tree found in 2012, this infestation was characterized as being spread out and heavy (NM, personal observation). No new infestations were reported in Ontario in 2020 but in 2021, *A. tsugae* was detected in Fort Erie from a posting made by a user on the iNaturalist website (josbees 2021) leading to additional delimitation and containment measures. In 2022, *A. tsugae* was detected near Pelham, Ontario during routine CFIA *A. tsugae* detection surveys, and in the township of Alnwick/Haldimand, near Grafton, by Canadian Forest Service and University of Guelph researchers who noted evidence of *A. tsugae* on the bark of trees and on branches at eye level while conducting plot assessments of a *T. canadensis* stand (MG, personal observation). This detection of *A. tsugae* was confirmed by the CFIA in July of 2022. In 2023 populations of *A. tsugae* were found in Hamilton on the grounds of the Royal Botanical Garden by employees of that facility; in Haldimand County as a result of a community-science detection project (Hemlock Woolly Adelgid Monitoring Network 2023); and near Lincoln by the Ontario Ministry of Natural Resources. In early 2024 populations of *A. tsugae* were found in Port Colborne during a routine CFIA detection survey for spotted lanternfly, *Lycorma delicatula* (White) (Hemiptera: Fulgoridae).

The age of the populations discovered in Ontario since 2019 have not been determined. One of us (JGF) surveyed hemlock from the Niagara Gorge to Prescot on the north shore of Lake Ontario in 2008 and found no evidence of infestations in any of the sites that were examined. We also conducted visual assessments of the infestations near Wainfleet in 2019 (CJKM, JGF) and near Grafton in 2023 (MG) that indicated these populations were well established and had been present for several years before detection. These populations have also been able to survive severe winter conditions and persist (MacQuarrie et al. 2024)

### Quebec, New Brunswick and Prince Edward Island

No detections of *A. tsugae* have been recorded in Quebec, New Brunswick or Prince Edward Island as of 2023. The first recorded CFIA survey for *A. tsugae* in Canada was done in Saint-Ignace-De-Stanbridge, Quebec in 1998, and the province has been regularly surveyed since 2009 (Fig. 5). The province of New Brunswick has also been regularly surveyed since 2009, with surveys in Prince Edward Island beginning in 2019 (Fig. 5). *Adelges tsugae* occurs in New York, southern Vermont and southern Maine and could be transported to New Brunswick and Quebec and from those infestations. Eastern hemlock in New Brunswick and Prince Edward Island would be at risk with spread of the adelgid from established populations in southern Nova Scotia (Fig. 3).

### Nova Scotia

The first detection in Nova Scotia of *A. tsugae* occurred in 2017 (Fig. 4A) from a report by an arborist that was confirmed by the CFIA. Following that initial detection, a visual ground survey was conducted at 131 sites by the CFIA, resulting in the detection of the pest in Annapolis, Digby, Yarmouth, Shelburne, and Queens counties (Fig. 5). Additional sites were inspected by Nova Scotia Department of Natural Resources and Renewables, Parks Canada, and non-governmental organizations. No new occurrences of *A. tsugae* were observed outside of these counties in 2018 and 2019. In 2020, *A. tsugae* was detected in Lunenburg County and an extensive survey was conducted at 87 sites across Nova Scotia in 2021, resulting in detections in Kings County. In 2022, CFIA conducted surveys at 66 sites, and no further infestations were identified. In 2023, new detections were made in Hants County and in Halifax County following reported sightings by a collaborating non-governmental organization and a homeowner, respectively that were confirmed by CFIA.

Observations of *A. tsugae* have been made year-round in Nova Scotia, with an apparent peak in July and August (Fig. 4B). Many of the records of observations in Nova Scotia in our combined database are drawn from iNaturalist records. In July and August progrediens generation ovisacs (i.e., sistens eggs) are typically at their largest size, and both generations of adults are conspicuous on foliage and so a peak in observations during those months may reflect their apparency to the public. This timing may also coincide to when more users are active and submitting reports to iNaturalist. A cursory inspection of the iNaturalist database also indicates that a peak of observations for *A. tsugae* also occurs in North America during this period (See seasonality map at: https://inaturalist.ca/taxa/61513-Adelges-tsugae).

### Range expansion in Canada

We estimated an expansion rate of 12.6 ± 8.2 km/year in the distribution of *A. tsugae* in Nova Scotia based on the change in the position of the population’s centroid with a northward expansion rate of 20.5 ± 27.21 km/year. The large variability in the northward expansion rate can be partly attributed the presence of a new detection of the adelgid in 2021 that was 76.5 Km from any known populations in 2020. When we do not include that 2021 detection in our calculations the rate of range expansion is 13.4 ± 12.6 Km/year. These estimates are consistent with observations of range expansion in the United States of America, over longer time scales (Morin et al. 2009; Turner et al. 2011; Fitzpatrick et al. 2012). Northward expansion in Nova Scotia also does not appear to have been limited by overwintering conditions (MacQuarrie et al. 2024). In Nova Scotia two infestations also appear to be correlated with human movement of infested material (JGO, personal observation).

We lack sufficient information to calculate a similar rate of expansion for Ontario, but there the dispersal of the insect may be limited due to forest fragmentation. In the Niagara region, and in many other parts of southern Ontario *T. canadensis* persists in protected areas and small woodlots (e.g., within Conservation Areas and parks) embedded within a matrix of urban areas and farmland (Shi 2024). Dispersal through this matrix is likely more difficult than in the eastern United States of America and Nova Scotia where forests are more contiguous. This may change, however, should *A. tsugae* infest more of the forests north of Lake Ontario, where *T. canadensis* is more common and forest conditions are more similar to those other invaded areas.

#### Summary and Conclusion

*Adelges tsugae* is perhaps the only species in Canada that has distinct native and invasive populations. We have summarized the present state of knowledge on the distribution of *A. tsugae* in Canada. In British Columbia the distribution of the adelgid is well characterized in the lower mainland and on Vancouver Island but is poorly surveyed in the interior and northern Pacific coast. These under-surveyed areas could host populations of the insect and natural enemies that may be better-matched to the climate in eastern Canada (Mausel et al. 2011), though populations of *L. nigrinus* from coastal British Columbia appear to survive winters in Nova Scotia (JGF, personal observation). This assumes, though, that these populations of natural enemies can be found and exploited. In western North America outbreaks of *A. tsugae* are usually ephemeral, lasting only one or two years which makes locating and exploiting such populations as a source of natural enemies difficult. Though this difficulty is mitigated somewhat in urban settings, where it is warm and lush and populations are more easy to survey and exploit. In Washington state *A. tsugae* is regulated by natural enemies on both native *T. heterophylla* and introduced *T. canadensis* (Crandall et al. 2022) but we lack an understanding of the ecology of *A. tsugae* in its native range in British Columbia, aside from the role of *L. nigrinus* in regulating populations (Zilahi-Balogh et al. 2003; Zilahi-Balogh et al. 2012). This includes the relationship between *A. tsugae* and *T. mertensiana* which has not been explored and could prove informative for understanding the host-herbivore dynamics in this system and suggest solutions for developing resistance in *T. canadensis*.

In eastern Canada *A. tsugae* continues to expand its range in Ontario and Nova Scotia. There are probably no climatic boundaries preventing the insect from infesting most of the maritime provinces (Kantola et al. 2019; MacQuarrie et al. 2024; Cornelson et al. in press), suggesting *T. canadensis* populations in New Brunswick and Prince Edward Island are at similar risk from the adelgid. The climate in Ontario, however, may prevent the insect from infesting the entire range of hemlock in that province, but those climate barriers are likely to disappear under existing climate change projections (Cornelson et al. in press). This is supported by recent detections of the insect in the northern part of Michigan’s lower peninsula where climate conditions are similar to that of central Ontario. In eastern Canada it will be important to understand and develop tools to mitigate the impacts on stand structure and composition (Parker et al. 2023), and determine the pathways of invasion into and within Canada to optimize future survey efforts (Yemshanov et al. 2020) and management options.

## Acknowledgements

We thank R Fournier (Natural Resources Canada) for assistance with GIS analyses, the technicians and researchers of the Forest Insect and Disease Survey, Canadian Food Inspection Agency and other agencies, and the users of the iNaturalist for contributing records of *A. tsugae*. This work was supported by Natural Resources Canada and the Canadian Food Inspection Agency.

## Literature Cited

British Columbia Ministry of Forests - Research Branch. 1996. The ecology of the interior Cedar-Hemlock zone.

British Columbia Ministry of Forests - Research Branch. 1997. The ecology of the mountain hemlock zone.

British Columbia Ministry of Forests - Research Branch. 1998. The ecology of the engelmann spruce- subalpine fir zone.

Canadian Food Inspection Agency. 2012. Plant Protection Survey Report 2011-2012. Canadian Food Inspection Agency, Ottawa, Ontario.

Canadian Food Inspection Agency. 2013. Plant Protection Survey Report 2012-2013. Canadian Food Inspection Agency, Ottawa, Ontario.

Canadian Food Inspection Agency. 2014. Plant Protection Survey Report 2013-2014. Canadian Food Inspection Agency, Ottawa, Ontario.

Canadian Food Inspection Agency. 2015. Plant Protection Survey Report 2014-2015. Canadian Food Inspection Agency, Ottawa, Ontario.

Canadian Food Inspection Agency. 2016. Plant Protection Survey Report 2015-2016. Canadian Food Inspection Agency, Ottawa, Ontario.

Canadian Food Inspection Agency. 2017. Plant Protection Survey Report 2016-2017. Canadian Food Inspection Agency, Ottawa, Ontario.

Canadian Food Inspection Agency. 2018. 2017-2018 Plant Protection Survey Report. Canadian Food Inspection Agency, Ottawa, Ontario.

Canadian Food Inspection Agency. 2019. 2018-2019 Plant Protection Survey Report. Canadian Food Inspection Agency, Ottawa, Ontario.

Canadian Food Inspection Agency. 2020. April 1, 2019 to March 31, 2020 Plant Health Survey Report. Canadian Food Inspection Agency, Ottawa, Ontario.

Canadian Food Inspection Agency. 2021. April 1, 2020 to March 31, 2021 Plant Health Survey Report. Canadian Food Inspection Agency, Ottawa, Ontario.

Canadian Food Inspection Agency. 2022. April 1, 2021 to March 31, 2022 Plant Health Survey Report. Canadian Food Inspection Agency, Ottawa, Ontario.

Canadian Food Inspection Agency. 2023. April 1, 2022 to March 31, 2023 Plant Health Survey Report. Canadian Food Inspection Agency, Ottawa, Ontario.

Chystal, R.N. 1916. The forest insect problem in Stanley Park. Proceedings of the Entomological Society of British Columbia, 9: 63–66.

Cornelson, C., MacQuarrie, C.J.K., and Lee, S.-I. in press. Modeling the distribution of hemlock woolly adelgid under several climate change scenarios. Canadian Journal of Forest Research.

Crandall, R.S., Lombardo, J.A., and Elkinton, J.S. 2022. Top-down regulation of hemlock woolly adelgid (Adelges tsugae) in its native range in the Pacific Northwest of North America. Oecologia, 199: 599–609. 10.1007/s00442-022-05214-8.

Elkinton, J.S., Lombardo, J.A., Roehrig, A.D., McAvoy, T.J., Mayfield, A., and Whitmore, M. 2017. Induction of cold hardiness in an invasive herbivore: The case of hemlock woolly adelgid (Hemiptera: Adelgidae). Environmental Entomology, 46: 118–124. 10.1093/ee/nvw143.

Ellison, A.M., Orwig, D.A., Fitzpatrick, M.C., and Preisser, E.L. 2018. The past, present, and future of the hemlock woolly adelgid (Adelges tsugae) and Its ecological interactions with eastern hemlock (Tsuga canadensis) forests. Insects, 9: 172. 10.3390/insects9040172.

Emilson, C., Bullas-Appleton, E., McPhee, D., Ryan, K., Statsny, M., Whitmore, M., et al. 2018. Hemlock woolly adelgid management plan for Canada, Natural Resources Canada Canadian Forest Service Information Report GLC-X-21. Sault Ste. Marie, Ontario, Canada.

Eschtruth, A.K., Evans, R.A., and Battles, J.J. 2013. Patterns and predictors of survival in Tsuga canadensis populations infested by the exotic pest Adelges tsugae: 20 years of monitoring. Forest Ecology and Management, 305: 195–203. 10.1016/j.foreco.2013.05.047.

Fidgen, J.G., Whitmore, M.C., MacQuarrie, C.J.K., and Turgeon, J.J. 2021. Detection of Adelges tsugae (Hemiptera: Adelgidae) wool using Velcro-covered balls. The Canadian Entomologist, 153: 640–650. 10.4039/tce.2021.24.

Fidgen, J.G., Whitmore, M.C., Studens, K.D., Macquarrie, C.J.K., and Turgeon, J.J. 2020. Sticky traps as an early detection tool for crawlers of Adelges tsugae (Hemiptera: Adelgidae). Journal of Economic Entomology, 113: 496–503. 10.1093/jee/toz257.

Fitzpatrick, M.C., Preisser, E.L., Porter, A., Elkinton, J., and Ellison, A.M. 2012. Modeling range dynamics in heterogeneous landscapes: Invasion of the hemlock woolly adelgid in eastern North America. Ecological Applications, 22: 472–486. 10.1890/11-0009.1.

Fryer, J.L. 2018. Tree species distribution maps from Little’s “Atlas of United States trees” series. In: Fire Effects Information System. U.S. Department of Agriculture, Forest Service, Rocky Mountain Research Station, Fire Sciences Laboratory (Producer). Available from: https://www.fs.usda.gov/database/feis/pdfs/Little/aa_SupportingFiles/LittleMaps.html [92602]. [accessed 1 February 2024].

Global Biodiversity Information Facility. 2023. GBIF occurrence download. Available from 10.15468/dl.je8tvc [accessed 27 July 2023].

Godman, R.M. and Lancaster, K. 1990. Tsuga canadensis (L.) Carr Eastern Hemlock. In Silvics of North America. Volume 1, Conifers. USDA Agriculture Handbook 271. Edited by R.M. Burns and B.H. Honkala. U.S. Department of Agriculture, Forest Service, Washington, DC. Pp. 604–612.

Goldstein, J., Park, J., Haran, M., Liebhold, A., and Bjornstad, O.N. 2019. Quantifying spatio-temporal variation of invasion spread. Proceedings of the Royal Society of London, Series B: Biological Sciences, 286: 20182294. 10.1098/rspb.2018.2294.

Havill, N.P., Montgomery, M.E., Yu, G., Shiyake, S., and Caccone, A. 2006. Mitochondrial DNA from hemlock woolly adelgid (Hemiptera: Adelgidae) suggests cryptic speciation and pinpoints the source of the introduction to eastern North America. Annals of the Entomological Society of America, 99: 195–203. 10.1603/0013-8746(2006)099[0195:Mdfhwa]2.0.Co;2.

Havill, N.P., Shiyake, S., Lamb Galloway, A., Foottit, R.G., Yu, G., Paradis, A., et al. 2016a. Ancient and modern colonization of North America by hemlock woolly adelgid, Adelges tsugae (Hemiptera: Adelgidae), an invasive insect from East Asia. Molecular Ecology, 25: 2065–2080. 10.1111/mec.13589.

Havill, N.P., Shiyake, S., Lamb Galloway, A., Foottit, R.G., Yu, G., Paradis, A., et al. 2016b. Ancient and modern colonization of North America by hemlock woolly adelgid, Adelges tsugae (Hemiptera: Adelgidae), an invasive insect from East Asia [Dataset]. Dryad. 10.5061/dryad.375dk. Hemlock Woolly Adelgid Monitoring Network. 2023. Available from https://www.invasivespeciescentre.ca/take-action/hemlock-woolly-adelgid-monitoring-network/ [accessed 20 Feb 2024].

Hemlock Woolly Adelgid National Initiative. 2024. Hemlock Woolly Adelgid National Initiative. Available from https://hemlock-woolly-adelgid-national-initiative-gmsts.hub.arcgis.com/ [accessed 9 July 2024].

Hopping, G.R. 1939. Forest insects of the season 1938 in British Columbia and Western Alberta. The Canadian Insect Pest Review, 17: 86–89.

josbees. 2021. iNaturalist observation. Available from https://inaturalist.ca/observations/75602865 [accessed 20 Feb 2024].

Kantola, T., Tracy, J.L., Lyytikäinen-Saarenmaa, P., Saarenmaa, H., Coulson, R.N., Trabucco, A., et al. 2019. Hemlock woolly adelgid niche models from the invasive eastern North American range with projections to native ranges and future climates. iForest - Biogeosciences and Forestry, 12: 149–159. 10.3832/ifor2883-012.

Lombardo, J.A. and Elkinton, J.S. 2017. Environmental adaptation in an asexual invasive insect. Ecology and Evolution, 7: 5123–5130. 10.1002/ece3.2894.

Loo, J. and Ives, N. 2003. The Acadian forest: Historical condition and human impacts. The Forestry Chronicle, 79: 462–474. 10.5558/tfc79462-3.

MacQuarrie, C.J.K., Derry, V., Gray, M., Mielewczyk, N., Crossland, D., Ogden, J.B., et al. 2024. Effect of a severe cold spell on overwintering survival of an invasive forest insect pest. Current Research in Insect Science, 5: 100077. 10.1016/j.cris.2024.100077.

Mausel, D.L., Van Driesche, R.G., and Elkinton, J.S. 2011. Comparative cold tolerance and climate matching of coastal and inland Laricobius nigrinus (Coleoptera: Derodontidae), a biological control agent of hemlock woolly adelgid. Biological Control, 58: 96–102. 10.1016/j.biocontrol.2011.04.004.

Mayfield III, A.E., Bittner, T.D., Dietschler, N.J., Elkinton, J.S., Havill, N.P., Keena, M.A., et al. 2023. Biological control of hemlock woolly adelgid in North America: History, status, and outlook. Biological Control, 185: 105308. 10.1016/j.biocontrol.2023.105308.

McAvoy, T., Régnière, J., St-Amant, R., Schneeberger, N., and Salom, S. 2017. Mortality and recovery of hemlock woolly adelgid (Adelges tsugae) in response to winter temperatures and predictions for the future. Forests, 8: 497. 10.3390/f8120497.

McClure, M.S. 1987. Biology and control of hemlock woolly adelgid. Bulletin 851. The Connecticut Agricultural Experiment Station, New Haven, Connecticut, United States of America.

McClure, M.S. 1989. Evidence of a polymorphic life cycle in the hemlock woolly adelgid, Adelges tsugae (Homoptera: Adelgidae). Annals of the Entomological Society of America, 82: 50–54. 10.1093/aesa/82.1.50.

McClure, M.S. 1990. Role of wind, birds, deer, and humans in the dispersal of hemlock woolly adelgid (Homoptera: Adelgidae). Environmental Entomology, 19: 36–43. 10.1093/ee/19.1.36.

McClure, M.S. 1991. Density-dependent feedback and population cycles in Adelges tsugae (Homoptera: Adelgidae) on Tsuga canadensis. Environmental Entomology, 20: 258–264. 10.1093/ee/20.1.258.

Means, J.E. 1990. Tsuga mertensiana (Bong.) Carr. Mountain Hemlock. In Silvics of North America. Volume 1, Conifers. USDA Agriculture Handbook 271. Edited by R.M. Burns and B.H. Honkala. U.S. Department of Agriculture, Forest Service, Washington, DC. Pp. 623–634.

Morin, R.S., Liebhold, A.M., and Gottschalk, K.W. 2009. Anisotropic spread of hemlock woolly adelgid in the eastern United States. Biological Invasions, 11: 2341–2350. 10.1007/s10530-008-9420-1.

Moss, E.H. and Packer, J.G. 1983. The Flora of Alberta, Second Edition. University of Toronto Press, Toronto, Ontario.

Natural Resources Canada Canadian Forest Service. 2023. FIAS Document Library. Available from https://www.exoticpests.gc.ca/documents [accessed 1 August 2023].

North American Plant Protection Organization. 2012. Detection and eradication of hemlock woolly adelgid (Adelges tsugae Annand) in Etobicoke, Ontario. Available from https://www.pestalerts.org/official-pest-report/detection-and-eradication-hemlock-woolly-adelgid-adelges-tsugae-annand [accessed 9 November 2020].

Nova Scotia Department of Lands & Forestry and Nova Scotia Department of Environment and Climate Change. 2024. Hemlock woolly adelgid pre-assessment risk analysis. Nova Scotia Department of Lands & Forestry – Fleet Services and Forest Protection Division, Risk Services Group and Nova Scotia Department of Environment and Climate Change – Protected Areas Branch.

Orwig, D.A. and Foster, D.R. 1998. Forest response to the introduced hemlock woolly adelgid in southern New England, USA. Journal of the Torrey Botanical Society, 125: 60–73. 10.2307/2997232.

Packee, E.C. 1990. Tsuga heterophylla (Raf.) Sarg. Western Hemlock. In Silvics of North America. Volume 1, Conifers. USDA Agriculture Handbook 271. Edited by R.M. Burns and B.H. Honkala. U.S. Department of Agriculture, Forest Service, Washington, DC. Pp. 613–622.

Paradis, A., Elkinton, J., Hayhoe, K., and Buonaccorsi, J. 2008. Role of winter temperature and climate change on the survival and future range expansion of the hemlock woolly adelgid (Adelges tsugae) in eastern North America. Mitigation and Adaptation Strategies for Global Change, 13: 541–554. 10.1007/s11027-007-9127-0.

Parker, W.C., Derry, V., Elliott, K.A., MacQuarrie, C.J.K., and Reed, S. 2023. Applying three decades of research to mitigate the impacts of hemlock woolly adelgid on Ontario’s forests. The Forestry Chronicle, 99: 205–225. 10.5558/tfc2023-024.

Plant Protection Act. 1990. Plant Protection Act, S.C. 1990, c. 22.

R Core Team. 2019. R: A language and environment for statistical computing. R Foundation for Statistical Computing, Vienna, Austria.

Ratnasingham, S. and Hebert, P.D. 2007. bold: The Barcode of Life Data System (http://www.barcodinglife.org). Molecular Ecology Notes, 7: 355–364. 10.1111/j.1471-8286.2007.01678.x.

Russo, N.J., Cheah, C.A.S.J., and Tingley, M.W. 2016. Experimental evidence for branch-to-bird transfer as a mechanism for avian dispersal of the hemlock woolly adelgid (Hemiptera: Adelgidae). Environmental Entomology, 45: 1107–1114. 10.1093/ee/nvw083.

Russo, N.J., Elphick, C.S., Havill, N.P., and Tingley, M.W. 2019. Spring bird migration as a dispersal mechanism for the hemlock woolly adelgid. Biological Invasions, 21: 1585–1599. 10.1007/s10530-019-01918-w.

Shi, Z. 2024. Identifying and classifying eastern hemlock (Tsuga canadensis) from surrounding species in Ontario forest using time series Sentinel-2 satellite imagery and phenological parameters, University of Guelph.

Turner, J.L., Fitzpatrick, M.C., and Preisser, E.L. 2011. Simulating the dispersal of hemlock woolly adelgid in the temperate forest understory. Entomologia Experimentalis et Applicata, 141: 216–223. 10.1111/j.1570-7458.2011.01184.x.

Van Sickle, A., Fiddick, R.L., and Wood, C.S. 2001. Forest insect and disease survey in the Pacific Region. Journal of the Entomological Society of British Columbia, 98: 169–176.

Venables, E.P. and Hopping, R. 1938. Untitled reports of Adelges tsugae. The Canadian Insect Pest Review, 16: 173.

Wallace, M.S. and Hain, F.P. 2000. Field surveys and evaluation of native and established predators of the hemlock woolly adelgid (Homoptera: Adelgidae) in the southeastern United States. Environmental Entomology, 29: 638–644. 10.1603/0046-225x-29.3.638.

Yemshanov, D., Haight, R.G., MacQuarrie, C.J.K., Koch, F.H., Liu, N., Venette, R., et al. 2020. Optimal planning of multi-day invasive species surveillance campaigns. Ecological Solutions and Evidence, 1: e12029. 10.1002/2688-8319.12029.

Zilahi-Balogh, G.M.G., Humble, L.M., Lamb, A.B., Salom, S.M., and Kok, L.T. 2012. Seasonal abundance and synchrony between Laricobius nigrinus (Coleoptera: Derodontidae) and its prey, the hemlock woolly adelgid (Hemiptera: Adelgidae). The Canadian Entomologist, 135: 103–115. 10.4039/n02-059.

Zilahi-Balogh, G.M.G., Salom, S.M., and Kok, L.T. 2003. Development and reproductive biology of Laricobius nigrinus, a potential biological control agent of Adelges tsugae. BioControl, 48: 293–306. 10.1023/a:1023613008271.

